# Kolmogorov-Smirnov and Shapiro-Wilk single distribution analysis methods in examining sample structure, strength of genetic background, and effects of interventions in lifespan and aging studies

**DOI:** 10.1101/2024.12.12.628153

**Authors:** Alexander V. Konopatov, Yuli V. Shidlovsky, Alexander A. Shtil, Oleg V. Bylino

**Affiliations:** Department of Gene Expression Regulation in Development, Institute of Gene Biology, Russian Academy of Sciences, 119334 Moscow, Russia; Blokhin National Medical Research Center of Oncology, 115522 Moscow, Russia; Department of Biology and General Genetics, Sechenov First Moscow State Medical University (Sechenov University), 119992 Moscow, Russia; Department of Regulation of Genetic Processes, Laboratory of Molecular Organization of the Genome, Institute of Gene Biology RAS, 119334 Moscow, Russia; Center for Precision Genome Editing and Genetic Technologies for Biomedicine, Institute of Gene Biology, Russian Academy of Sciences, 119334 Moscow, Russia

**Author notes:** Correspondence (O.V.B.); (A.A.S.). ‘I think it may fairly be assumed, in the light of what we now know, that no other measure will, statistically speaking, furnish so delicate and precise a measure of the general constitutional fitness of individuals as will their duration of life.’. Raymond Pearl, 1923.

## Abstract

The analysis of lifespan and ageing involves a variety of calculations based on survival data. One of these is the comparison of survival curves. However, inspection of survival curves provides little information about the frequency of phenotypes within samples. Here we propose to divide the distribution into intervals to obtain information about the sample structure. Then, we apply the normality criterion to the distributions using the Kolmogorov-Smirnov and Shapiro-Wilk tests. Using this approach, we reveal the destabilising effect of mutations on ontogenesis and lifespan, and estimate the strength of the genetic background of the lines. The proposed methodology allows effective estimation of sample structure and adds a new layer of information to longevity studies.

## Introduction

Lifespan is the most complex quantitative trait in humans. Surprisingly, survival patterns in Drosophila replicate those of humans. This phenomenon was remarkably discovered early in the twentieth century by the eminent researcher Professor Raymond Pearl (Pearl and Parker, 1922), who, on the basis of studies in Drosophila, proposed the use of lifespan as the main indicator of Darwinian fitness (Pearl, 1923). Although not all researchers share this view because, as postulated by evolutionary theory, the primary measure of a genotype’s fitness is the number of offsprings it leaves behind (Michod, 1986; Pfenninger, 2017), lifespan is nonetheless among the major components of Darwinian fitness/life-history traits of individuals, along with age of puberty, fecundity, fertility, and age-specific survival dynamics of individuals (Flatt, 2020). Thus, lifespan is, if not the most important, at least one of the most important components of fitness. Therefore, it is crucial to know the patterns of distribution of lifespan data of individuals in the samples studied, and to understand what shifts in this distribution depend on.

Studies of lifespan and aging are inherently linked to the construction of Kaplan-Meier survival curves (Flatt and Partridge, 2018). A survival curve reflects the attrition of the cohort (individuals born at the same time) or population being studied from time. Survival curves are common in clinical research, for example in recording the survival of cancer patients on chemotherapy, and in lifespan and ageing studies, recording the mortality of short-lived model organisms in response to the effects of pharmacological and genetic interventions (introduced mutations, transgenes, therapies administered). To assess the reliability of differences in survival curves, comparisons between samples are used based on the median 50th percentile for sample mortality), mean (arithmetic mean of the lifespan of all individuals in the sample) and maximum (90% sample mortality) lifespan. Appropriate statistical tests, such as the Kolmogorov-Smirnov two-sample test (Fleming et al., 1980) or the log-rank test (Mantel-Cox test) (Mantel, 1966), are common for estimating the former. The Wang-Allison test is used to compare samples by maximum lifespan (Wang et al., 2004). However, all these types of tests provide information only on the significance of differences between samples at certain points in time. Survival curves also provide only a general idea of how mortality increases over time in the cohort under study, while a clear picture of the distribution of phenotypes by lifespan within the cohort under study is not shown by the survival curve. This requires the development of new approaches to analysing survival data that clearly show differences in the structure of the samples being compared (problem 1).

Another important problem in studies of lifespan and aging is the lack of mathematical criteria for assessing the strength of the genetic background (genotype) of the studied lines, as well as the effect on lifespan of various interventions aimed at increasing it (problem 2). Lifespan, as mentioned above, is an integral indicator of the genotype health of a line and it is believed that if the cohort survival curve has a convex shape, then those animals have better genetic health than lines with linear or concave survival curves (Piper and Partridge, 2016). However, specific mathematical criteria for assessing the strength of the genetic background of lines based on lifespan data have not been developed. Some approximation of such an assessment is the construction of mortality curves described by the Gompertz equation, a mortality law formulated at the end of the first quarter of the 19th century by Benjamin Gompertz (Kirkwood, 2015). However, in the case of mortality curves, the genetic health of lines is assessed based on the numerical value of the equation parameter α (baseline vulnerability, initial/early mortality), whereas it is certainly more reliable to assess the health of lines based on the entire distribution of mortality data of a cohort over its entire lifespan period. Thus, the development of mathematical criteria for the integral assessment of the genetic health of lines is an important challenge, as it will allow estimation based on the lifespan of all individuals in the sample, not just those who died during certain periods.

Here we considered methods for comparing and analysing survival curves and proposed to break survival data into intervals to solve the problem of sample structure (problem #1). The analysis showed that using this approach, it is possible to visually assess the distributions of phenotype frequencies by life expectancy within a sample. Partitioning into intervals allowed a more efficient approach to problem 2. In order to assess the genetic background of the lines and the effect of interventions on ontogeny and longevity, the authors applied the criterion of normality of distribution to the survival data. They found that lines with reduced or increased longevity were characterised by abnormal distributions and/or large deviation from normality. Estimating the genetic background of lines using the normality criterion allows us to draw conclusions based on the totality of all data in the whole sample and not only based on early life mortality, as postulated by Gompertz’s law.

The use of the normality criterion may become an important tool in studies of lifespan and aging, help integrate estimates predicted by simpler methods, allow the selection and comparison of pairs of lines with similar genetic health, better understand and describe the processes occurring in populations, and help monitor the impact of genetic and pharmacological interventions on ontogenesis and lifespan.

## 1. Materials and methods

At least 40 tests have been developed to analyse the distribution in single samples (Arnastauskaitė et al., 2021). The two main criteria for a statistical test are convenience (ease of calculation) and demonstrativeness (ability to identify patterns). The most popular and widespread are the Kolmogorov-Smirnov test (KS-test) (Facchinetti, 2009; Simard and L’Ecuyer, 2011) and the Shapiro-Wilk test (SW-test) (Royston, 1993; Mara, 2011). Both tests in their classical versions fulfil both criteria.

Drosophila melanogaster lifespan (LS) series were used as data: 5,6,6,6,6,6,6,6,6,6,6,6,6,8,9,9,9,9,10 etc., where each number shows the LS of one individual fly specimen. 100% mortality of flies was observed between 57 and 82 day. Between 163 and 472 flies participated in the experiments, depending on the cohort observed. WT-like laboratory lines Canton S, Oregon RS and lines containing mutant alleles of the white gene — w^1118^ and w^67c23^ — were studied. The w^67c23^ allele was isolated from the chromosome of the standard laboratory line *y^1^ w*^67c23^ via recombination with chromosomes of wild-type lines. The *w^1118^* allele was taken from the #3605 Bloomington collection. The genetic background of the mutant lines was aligned by 10 backcrosses to the corresponding wild-type lines. LS data from two experiments conducted at different times, in 2024 (allele w^1118^) and in 2012 (w^67c23^), were used for analyses. The spring 2012 experiment was performed at room temperature (22-23 °C), while the spring 2024 experiment was performed at 25 °C. In each experiment, male and female flies of mutant lines (w^1118^, or w^67c23^) were separately analysed with respective control lines by gender or by mutation. The Canton S and Oregon RS lines originated from the Drosophila collection of the Institute of Molecular Genetics of the Russian Academy of Sciences (IMG RAS).

### 1.1. Calculation of the Kolmogorov-Smirnov test for single samples

The one-sample KS test is a universal test that can be used to evaluate any type of data (discrete — 1,2,3,4, etc., continuous — 1.1, 1.2, 1.3, etc.) for adherence to any distribution (uniform, normal, exponential, etc.). The KS test is based on the search for the supremum — the point of maximum deviation (difference) between the experimental and expected integral functions of the normal distribution (CDF, cumulative distribution function) for a given sample (Fig. 1A) (Kolmogoroff, 1933; Feller, 1948).

**Figure 1.**
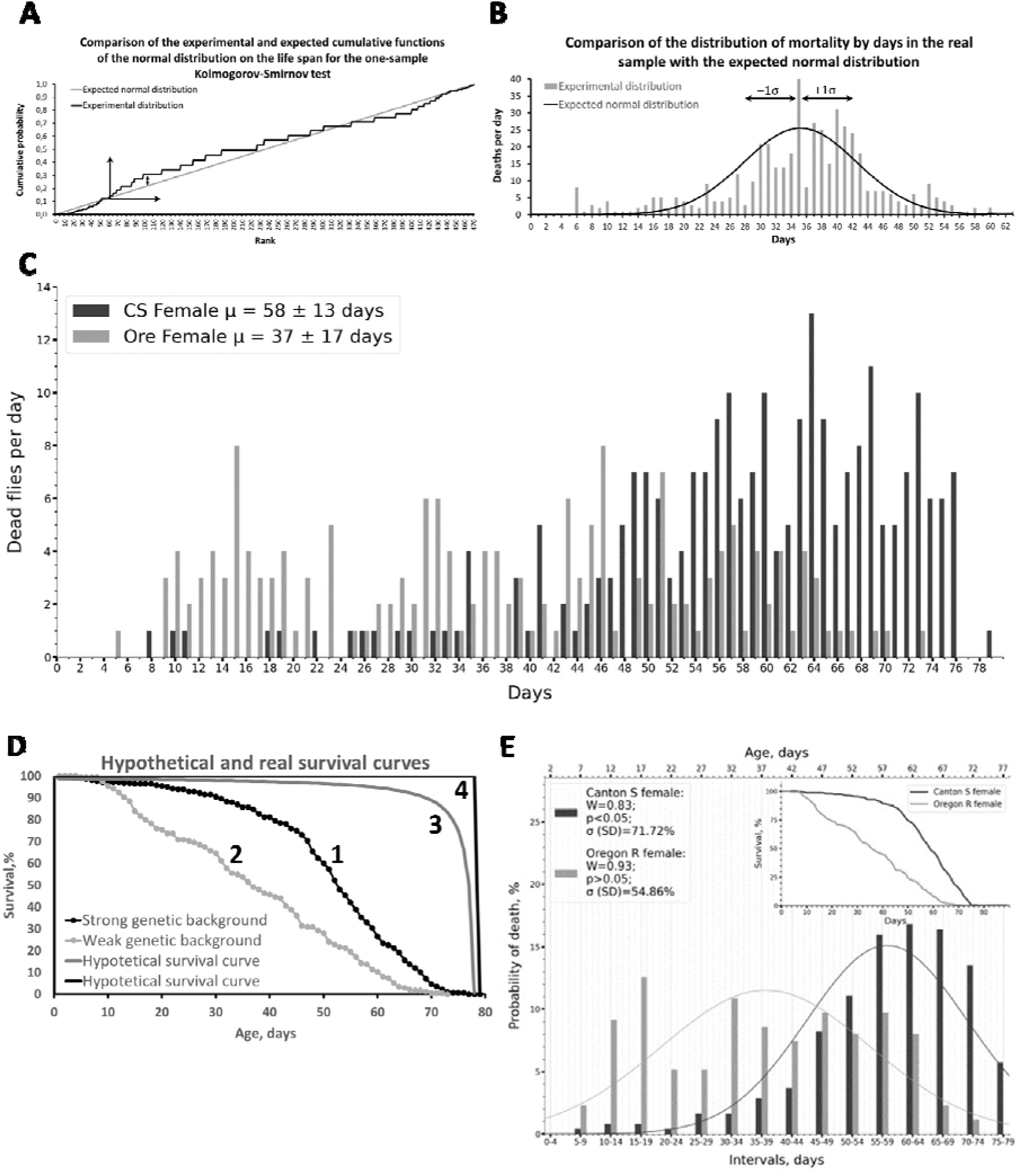
**(A, B)** Depicted are the principles of Kolmogorov-Smirnov (A) and Shapiro-Wilk (B) tests for analysing single distributions. **(А)**. The integral (cumulative) normal distribution function (CDF) tending to infinity (n→∞) is shown. The grey curve (straight line) shows the expected CDF at *n*→∞. The black curve shows the experimental CDF function constructed from the mean (µ) of the LS and the standard deviation (σ) of the study sample. The X-axis is the ranks (0,1,2,3, etc.) and the Y-axis is the integral of the mortality probability (probability density function). The arrow pointing upwards shows the increase in the experimental function due to the increase in the probability of mortality (more flies die in the next time interval than in the previous one). The arrow pointing to the right shows deviation due to repeated values of mortality (several individuals live the same number of days). If the curve deviates upwards, there is a succession of days with low mortality and vice versa — if the curve deviates downwards, then there is a succession of days with high mortality. The bidirectional arrow shows the supremum (maximum difference between the experimental and theoretical expected CDF). The size of the step of the graph (length) is directly proportional to the number of deaths per day. LS data from male males of the Oregon RS line (experiment with the allele^w1118^) were used. **(B)** The number of deaths per day for the real sample is shown (bars), as well as the normal distribution of the data (bell-shaped black curve) derived from µ (mean LS of the sample) and σ (standard deviation of µ) calculated from the LS of all individuals in the sample. The Shapiro-Wilk test compares the experimental distribution with a reference normal distribution. The calculation of the parameters of the normal distribution is given in the text in Section 3.2. ±1σ shows the limits of σ. The X-axis is days (0,1,2,3, etc.) and the Y-axis is the number of dead flies per day. Fly mortality data as in (A) are used. (C) Daily mortality data are presented for females of two different WT-like laboratory Drosophila lines with different genetic backgrounds, Canton S (black bars) and Oregon RC (grey bars) (data from an experiment with the allele #1). X-axis — days (0,1,2,3, etc.), Y-axis — number of dead flies per day. µ is the mean LS±σ in days. (D) Experimental survival curves of lines from (B) are shown (curve #1 — Canton S, #2 — Oregon RS), as well as hypothetical survival curves (#3,4) (details in text). (E) Experimental data distributions (bars) with phenotype frequencies obtained by partitioning samples from (C, D) into 5-day intervals are shown. Each bar shows the probability of fly death in 5 contiguous days. The bell-shaped black curves show the normal distributions calculated from the original sample data on LS. The calculation of normal distribution parameters is presented in section 3.2. W, p are the statistic and p-value of the Shapiro-Wilk test, respectively. The presented W and p values are obtained from the estimation of the distribution divided into intervals. µ and σ are calculated as for (B). For (E) The inset on the right shows survival curves for these samples.

CDF was found by the formula:

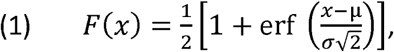

where x is the individual fly, µ is the mean, σ is the standard deviation (square root of the variance), erf is the error function:

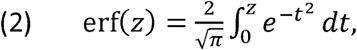

where 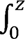 - is the integral from 0 to z, π is the number Pi, dt is the infinitely small increment of *t*, z in this case:

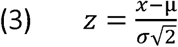

Differences between the experimental (black step curve) and expected (grey line) CDFs were found by subtracting the expected CDF from the experimental CDF at each point along the graph (see Figure 2). If the black step curve deviated upwards from the grey straight line, the difference was positive; if downwards, it was negative. The largest value of modulus of these differences is the supremum, the statistic of the KS test (D_n_).

**Figure 2:**
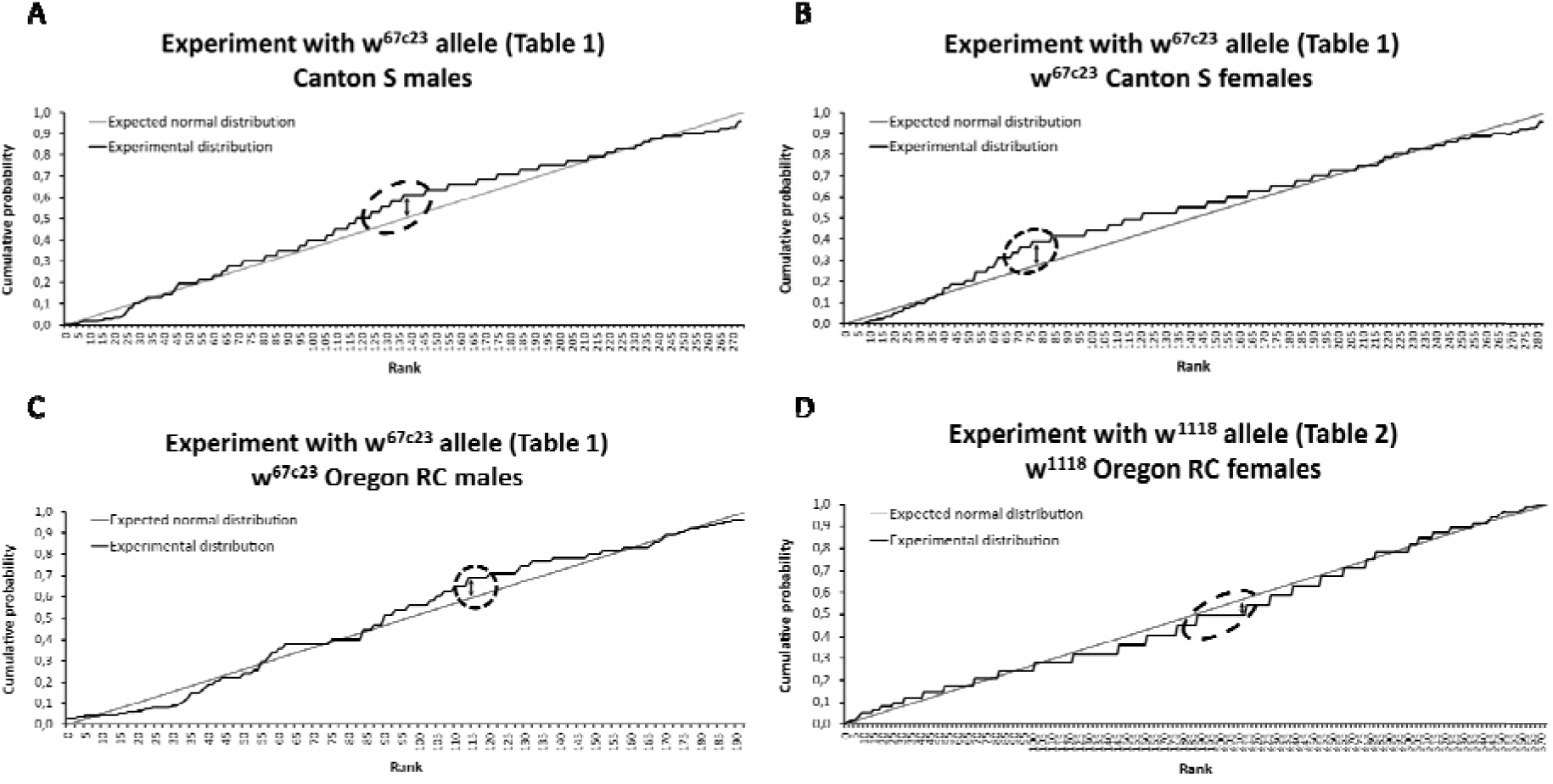
Shown are cases of maximum deviation of the supremum of the Kolmogorov-Smirnov function from the rank line (normal distribution function for the ranked expected sample with gradual increase from 1 to n in steps of 1). The cases from A to D correspond in order to the cases with normality violation, highlighted in grey in Tables 1,2.

The assessment of whether the distribution is normal is made in the KS test by comparing the value of the calculated test statistic D_n_ (D-experimental) with the D-critical value calculated for sample size *n* at the selected significance threshold α (α — 0.05, 0.01, etc.). Critical values for sample sizes <35 are extracted from Smirnov’s table (Smirnov, 1939) and for samples between 35 and 100 from Miller’s table (Miller, 1956). Also, for values >35 the formula d_α_/√N where d_α_ =1.36 at α=0.05. Other values for other significance levels are given in the works (Massey, 1951; Facchinetti, 2009). If D_n_ turns out to be less than D-critical — the distribution is considered normal. If the exact sample size is not available in the table, an alternative way to test the H0 (null) hypothesis (that the distribution is normal) is to find the p-value and compare it to the chosen significance threshold α (e.g., 0.05). In this case, if the p-value is greater than α, the distribution is considered normal. The calculation of the KS-test was carried out according to the generally accepted methodology (Feller, 1948; Massey, 1951).

The Kolmogorov formula was used to calculate the p-value (Kolmogoroff, 1933; Feller, 1948; Smirnov, 1948) which is convenient for both small samples (<35) and large samples:

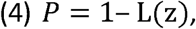

Where L(z):

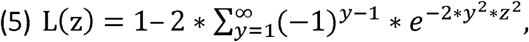

where 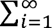 is the sum from 1 to ∞ (in practice, the sum of the first 10 y is significant, the rest do not add precision to the measurement), e is the exponent, 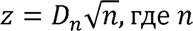 – sample size.

To calculate the p-value together with the classical Kolmogorov formula, other formulas have been proposed in the literature, for example, the formula proposed by Marsaglia (Marsaglia et al., 2003):

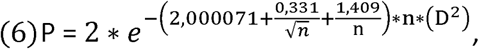

where, P is the p-value, e is the exponent, n is the sample size, and D is the statistic of the KC test.

We tested this formula and compared its performance with the classical Kolmogorov formula on a sample of random numbers, normally distributed data and our samples and concluded that the divergence of p-values occurs in the third digit. Hence, in terms of accuracy in predicting patterns, the Marsaglia formula differs weakly from the Kolmogorov formula. Importantly, the Marsaglia formula gives p-values greater than 1 (e.g., 1.8) in the case of normally distributed data (between 35 and 100 sample values). Therefore, we do not recommend using the Marsaglia formula to calculate p-value for the purposes of LS assessment.

### 1.2. Calculation of the Shapiro-Wilk test

The SW test, in contrast, assesses the symmetry of the data distribution by comparing the left and right sides of the distribution (Figure 1B) (Shapiro and Wilk, 1965). This test has a table of critical values against which the found value of the W test statistic is compared. The disadvantage of the standard (generalised) SW test is that both the table of critical values and the table of coefficients for calculating W are calculated only for small samples (≤50) (Shapiro and Wilk, 1965). However, LS studies involve analysing a much larger data set. Therefore, for sample sizes >50, the method of Royston, who proposed a modification of the formula for calculating W, the coefficients, and an algorithm for calculating p-value for large samples, can be used (Royston, 1993)

The Royston method uses a more sophisticated method of calculation than the standard (generalised) SW test, using a complex polynomial function. Unlike the (generalised) SW test, the Royston procedure for calculating W is the same for all sample sizes from 12 to 5000, but it proved to be too complex to satisfy the ease of use criterion. Therefore, although the Royston method is widely used in online calculators, the cumbersome calculation system and the risk of technical errors preclude the use of this method when analysing LS data thoroughly “manually”. In addition, when we performed calculations with this method (W and P), we found that all analysed distributions of CV data turned out to be non-normal (see Table 1,2 columns 6,7). Thus, the SW test as modified by Royston to calculate W was not indicative for large samples. Therefore, for the purposes of analysing samples containing a small number of values (<50), we propose to use the conventional (generalised) SW-test, combining its use with the procedure of dividing the sample (survival data) into intervals (see below). This reduces the sample size, greatly simplifies the calculation system, and allows us to approximate the p-value using Table #6 from the original paper by Shapiro and Wilk. (Shapiro and Wilk, 1965).

**Table 1.**
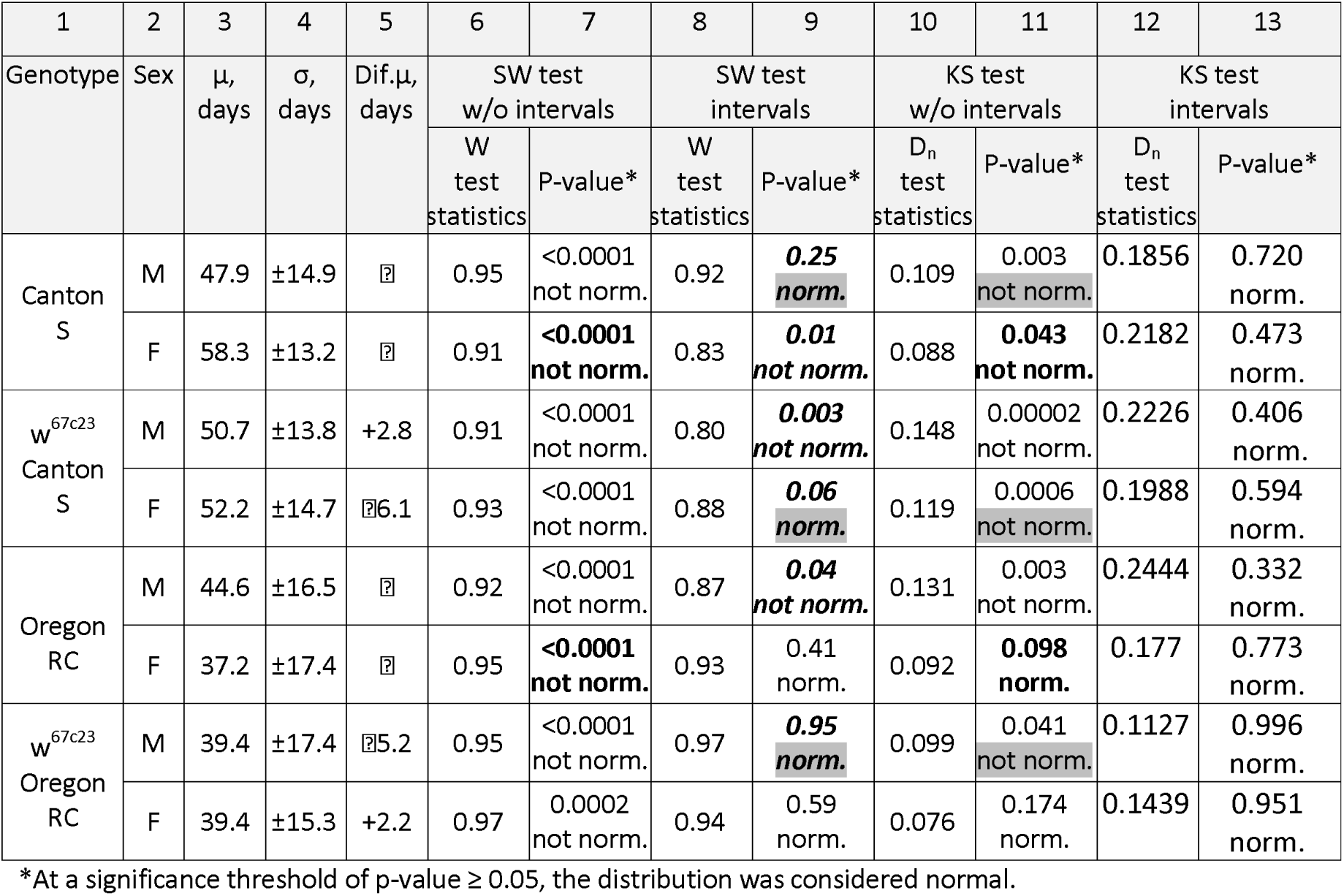
An experiment to measure LS with the w^67c23^ allele.

Shapiro and Wilk in Table #6 published W values for specific percentage points of P (0.01, 0.02, 0.05, 0.10, 0.50, 0.90, 0.95, 0.98, 0.99) for samples of size (n) 3 to 50 (Shapiro and Wilk, 1965). The values of the percentage points were found by Johnson’s approximation method using the S_B_ model, which fits the observed distribution to a normal distribution (Johnson, 1949). Whereas Royston proposed a method of finding the p-value (P, percentage point) for any W and almost any n without using a percentage point table (Royston, 1993)

In this paper, we used and present the exact p-value calculation algorithm from Royston’s paper, which is missing from the original Shapiro-Wilk paper (Royston, 1993).

The general final formula for calculating the p-value is as follows:

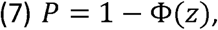

where Φ(*z*) is the function of the standard (µ = 0, σ = 1) integral normal distribution from z, µ is the arithmetic mean, σ is the standard deviation.

Φ(z) can be found as follows:

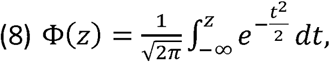

where z (z-score/standard score) — shows the deviation of the value of X from µ expressed as a quantity σ, *dt* is the infinitesimal increment of *t*.

z can be found as follows:

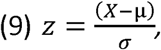

The value of Φ(*z*) can be calculated using the formula above, but also can be found in the z-score table (Warne, 2020) or by calculating the normal distribution formula (formula #19), for example, using the MS Excel function NORM.DIST(X; µ; σ; truth), where truth is cumulative (integral) function of the standard normal distribution.

Similar to Johnson, Royston’s work involves transforming the observed z distribution to a normal distribution:

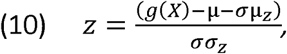

Where g(X) is the function of transforming the distribution of the original data to a normal distribution, µ_z_ is the transformed µ, σ_z_ is the transformed σ. Since here µ = 0, аnd σ = 1, we obtain:

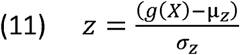

For sample sizes <12 and 12 to 2000, the formulas and coefficients for finding g(X), µ_z_ and σ_z_ take a different form and can be found in Table #1 of Royston’s original paper (Royston, 1993). Below we give the formulas and coefficients for finding g(X), µ_z_ and σ_z_ for samples from 12 to 2000:

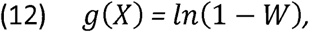

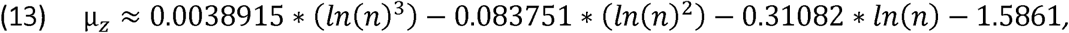

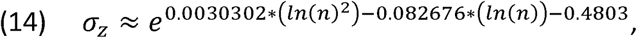

where, n is the sample size, e is the exponent, and W is the SW test statistic:

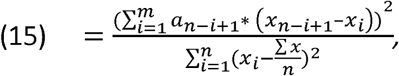

where, *x is* the sample (an ascending sorted series of numbers denoting the number of individuals who died in each interval), *n* is the sample size (how many numbers in the above series), m is the 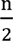 for even *n* and 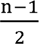 for odd 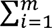 - the sum operator (index *i* takes values from 1 to m); *x i* is the row (sample) count term (1, 2, 3…m), *and а n*–*i+1* are the coefficients from Table #5 for calculating the value of the SW test statistic from the original paper (Shapiro and Wilk, 1965) The numerator of W is proportional to the square of the minimum variance of the linear standard deviation estimate σ, and the denominator is the sum of the squares of the variances with respect to the sample mean µ (sampling variance) (Royston, 1995).

### 1.3. Pearson correlation calculation

The Pearson correlation coefficient was calculated according to the standard methodology (Pearson, 1895) using the formula:

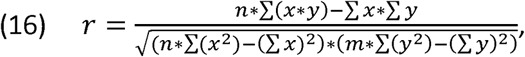

where n is sample size #1, m is sample size #2, x is the series of all values of sample #1, y is the series of all values of sample #2.

## 2. Results and discussion

Figure 1C shows a plot of the distribution of mortality by day for the two Drosophila lines we obtained, and Figure 1D shows the survival curves constructed from these data (curves #1, 2). Curves #3, 4 correspond to two hypothetical survival curves showing near-perfect (#3) and perfect (#4) cases of the distribution of LS data according to Flatt and Partridge (Flatt and Partridge, 2018). Curve #4 shows no mortality over the entire LS period of the cohort and simultaneous death of all individuals on the same day. This is a case of maximum rectangularisation of survival curves. Curve #3 shows linear mortality of the cohort over almost the entire LS period and an exponential increase in mortality at the end of life. Consistent with this, experimental curves #1 and #2 show different curvature and are characterised by a strong and weak genetic background, respectively. Mortality curves constructed from these data are characterised by diametrically opposite parameters of mortality accumulation according to the Gompertz equation:

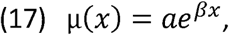

where, µ(x) — dynamics of mortality increase depending on age (x), α — constant 1 (baseline vulnerability of a cohort, line, population), β — constant 2 (exponent, aging rate, mortality accumulation rate).

Sample #1 is characterised by the following parameters: α = 0.2517, β = 0.5342, sample #2: α = 2.0672, β = 0.3832. Thus, the LS data of these two lines are an ideal model for comparison on the strength of genetic background.

### 2.1. Partitioning the survival data into intervals allows for a detailed assessment of the sampling structure

The application of the KS and SW tests to the LS series for the samples from Figure 1C

(at daily mortality) gave contradictory results (Table 1, Straight bold, columns 6,7,10,11). Thus, the SW test as modified by Royston showed a non-normal distribution in both cases, whereas the KS test showed a normal distribution in one case and a non-normal distribution in the other. However, a close look at the data in Figure 1C reveals that the data from sample 1 (light grey bars) resembles a uniform distribution, while the data from sample 2 (dark grey bars) resembles a bell-shaped distribution skewed towards the right edge. Thus, it is possible that at least one of the two distributions may be normal.

These results prompted us to analyse why distributions may be perceived as abnormal by the SW test. We noticed that the data from the two samples contain mortality dips (days with no mortality at all) and, on the contrary, outliers (days with mortality spikes). Thus, the problem of fluctuations in daily mortality is clearly identified. Such fluctuations will have a strong negative effect when assessing normality. This may be especially true for the KS test, where the final estimate is based on the deviation of a single point (supremum).

Accordingly, it was decided to pool the mortality of adjacent days by dividing the sample data into intervals. In addition to the fluctuations in daily mortality, splitting samples into intervals helps to address several other problems: i) imprecision in determining the date of death. For example, a fly may die immediately after inspection, on the same day, but will be recorded as dead the next day. ii) The stochastic effect of mobilising flies to a new feed. For example, if the feed spoils prematurely, the feed is replaced earlier rather than after 3 days, which may have a stochastic effect on LS. Obviously, dividing the data into intervals will offset the above “noise” and allow a more correct assessment of the distribution type. In addition, it will help to overcome the sample size limitation of 50 and use the simplified system of calculations from the original Shapiro-Wilk paper. The pooling procedure is essentially the same as counting dead flies every few days or even once a week, which is used in Drosophila LS experiments.

The classic approach to determining the size (class) of an interval is the rule of thumb proposed by Herbert Sturges (Sturges, 1926). Sturgess’ rule of thumb offers the following formula for finding the class of an interval:

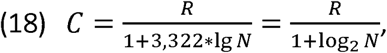

where, C is the size (class) of the interval (in our case, the number of contiguous days we combine); R is the difference between the maximum and minimum values of the sample (in our case, the maximum and minimum LS); N is the sample size (the number of flies whose LS distribution we determined); lgN is the decimal logarithm of N; log_2_ N is the logarithm of N on base 2 (equivalent to 3.322*lgN).

We performed the proposed procedure for our data with subsequent rounding to integers (according to rounding rules) and obtained the following results of interval sizes: 7, 7, 6, 5, 5, 7, 7, 6, 5, 7, 7, 7, 8, 8, 8, 8, 7, 7 (we used 16 samples from Table 1,2). However, in order to correctly compare the samples and their distributions, it is necessary to bring the data to the same form (use the same interval size). According to Sturgess the most “convenient” intervals are 2,5,10,20,50,100,200 etc. and the nearest “convenient” interval should be used. We settled on an interval length of 5 days, testing a range of 3—7 days. The analysis showed that it was the size of the interval of 5 days that provided acceptable visibility and detail of the graph while maintaining sufficient difference in the values of the W test statistic between the experimental sample and the control sample when applying the SW-test to them. The seemingly convenient value of 7 days created an oversimplified, sparse picture of the graph.

The results of partitioning into 5-day intervals of the two experimental samples shown in Figure 1C,D are presented in Figure 1E. We used a partitioning of the form: day 0-4, day 5-9, day 10-14, etc. It can be seen that the data distributions with 79 and 73 units (Figure 1C) are transformed into distributions with 15 and 14 units, respectively (Figure 1E). With the maximum LS of fly lines in the range of 57-82 days, the maximum number of mortality segments/intervals analysed would be 12-17.

Mortality in graphs can be expressed both in absolute numbers (number of individuals) and in relative units (probability, %). Comparison in relative units allows normalising the size of the samples being compared and is the most informative, as it allows comparison between data from experiments with different sample sizes.

It is important that when graphs are plotted, if one of the samples had a larger number of intervals (differed greatly in maximum LS from the other sample), the missing number of intervals with zero probability of mortality is added to the sample with a shorter range of LS. This makes it convenient to compare the obtained probability series on the graphs against each other. At the same time, when calculating further statistics of the SW and KS tests, all zero intervals (both early and late) available for a given sample are not considered in the calculation, as they may distort the values of the test statistics and their p-values.

Thus, the distributions obtained after dividing the sample into 5-day intervals clearly demonstrate the differences in phenotype frequencies (mortality probabilities) at different ages. Comparison of the resulting data is informative and allows us to effectively identify differences in sample structure, as well as to compare different samples with each other.

### 2.2. Normality criteria can be used when analysing survival data divided into intervals

To visually and formally describe the obtained interval distributions (bars), we superimposed the corresponding normal distribution (ND) on them (indicated by bell-shaped curves in Fig. 1E and following). The centre of the ND was taken as the mean LS of the original sample (not divided into intervals). For clarity of presentation on the graphs, the obtained ND values were multiplied by the number of individuals in the sample. In this case, the area under the graph was not equal to 1, but to the number of individuals in the corresponding sample and the graph correlated well with the size of the bars (for Fig. 1B, only the sample size was multiplied). The resulting ND values were then divided by the interval size (five) (for Fig. 1E and following). Without dividing by the interval size, the LS plot correlated poorly with the size of the bars and far exceeded their size.

The normal distribution for each day was calculated using the formula of the probability distribution function (probability distribution function, PDF):

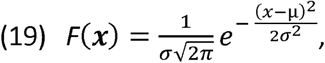

where σ is the standard deviation for the sample taken, *σ* is the distribution variance, *e* is the exponent, *µ* is the mean LS for the sample taken (mean lifespan, meanLS), *π* is the Pi number (3.14), *x* is the day. It is possible to calculate the normal distribution function, for example, using the MS Excel function NORM.DIST(X; µ; σ FALSE), where false is the PDF function, x is the day.

The mean LS of the original sample (µ) was calculated as:

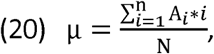

where 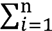 is the sum from the first to the last day of measurement (*n*), A_i_ is the number of flies which are dead on day *i*, N is the total number of flies in the sample, *i* is the day (1,2,3…). The standard deviation was taken as the variation of the mean LS, which was calculated according to the formula:

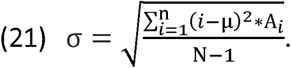

Thus, there was a clear opportunity to visually compare the parameters of the obtained interval distributions of phenotype frequencies by LS (mortality probabilities) with their normal distributions.

### 2.3. Use of Kolmogorov-Smirnov and Shapiro-Wilk normality estimation tests in describing the parameters of the survival distribution

Next, we described the obtained interval and initial distributions (not broken into intervals) by means of SW and KS tests for 16 samples (Table 1,2). We determined test statistics (numerical quantitative description) and normality parameters (qualitative description by the type “is” or “is not”).

It was found that the KS-test, when divided into intervals, qualifies all the 16 samples as normal, while the SW-test, on the contrary, qualifies all the not divided into intervals 16 samples as non-normal. Hence, in both cases, the test performance criterion is violated. Both cases were excluded from the analysis and were not considered further.

Thus, divided LS data into intervals is suitable for analysis by the SW test, while the not divided data is suitable for analysis by the KS test. We compared the concordance between the findings of the KS and SW tests in these cases. We found that in 4 cases out of 16 there was a mismatch of findings: when the SW test showed normality, the KS test showed a non-normal distribution (Table 1,2, cases of mismatch between the two tests are marked with grey shading). In order to find out why this happens, we took a deeper look at the performance of the KS-test in all 4 cases.

The analysis showed that the estimation of normality of the distribution using the KS test is, in general, strongly influenced by fluctuations in daily mortality (fluctuations in the structure of mortality data). Thus, we observed that the point corresponding to the supremum: i) can fall on a day with high mortality preceded by several days with low mortality (Figure 2 A, B, cases from Table 1 Canton S males and w^67c23^ Canton S females). In this case there is a strong shift of the function to the right (the difference between experimental and control CDF is positive), ii) can fall on a day with high mortality preceded by a day with minimal mortality (death of one individual) — there is a sharp/double increase in the height of the “step” of the function upwards and its strong deviation to the right (Fig. 2C, case from Table 1 w^67c23^ Oregon RC females) (difference is positive), iii) may fall on a day with very high mortality — there is a very strong deviation of the function to the right (Fig. 2D, case from Table 2 w^1118^ Oregon RC females) (the difference is negative, especially if the day of maximum mortality falls in the period of high mortality and the middle of the graph as in Figure 2D).

**Table 2:**
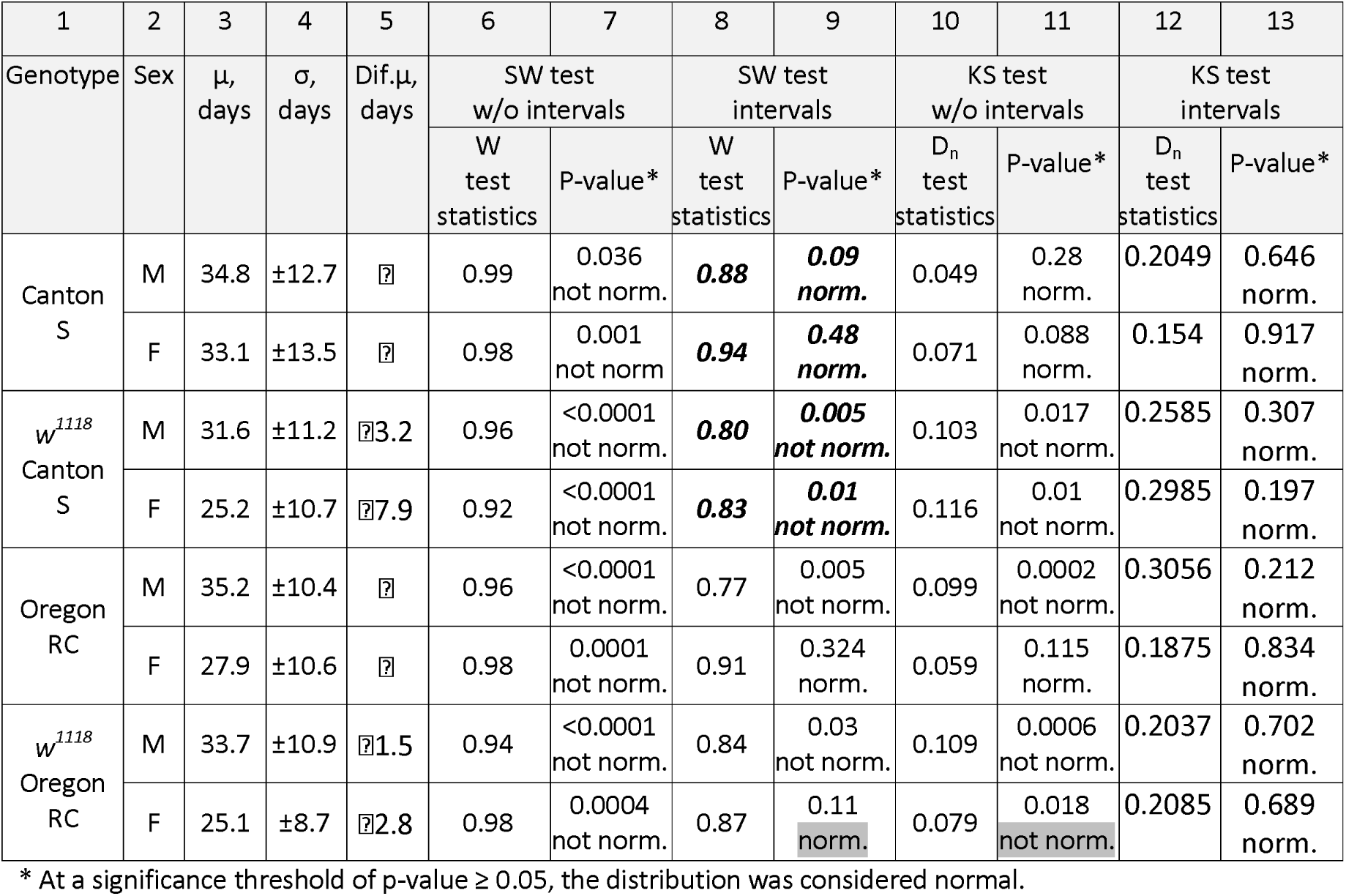
An experiment to measure LS with the w^67c23^ allele.

Thus, in all 4 cases considered, the KS-test defines the distribution as non-normal on the basis of violation of smoothness of variance of LS of one or another expression. Consequently, the principle of determining normality inherent in the KS-test is not insufficiently suitable for such type of data as LS and survival curve data, when there is a high fluctuation of mortality by days.

To better understand the regularities of the KS test, we conducted additional experiments. An interesting observation turned out to be that fluctuations of the mortality rate falling in the middle of the distribution, in the interval ±1σ, have the strongest effect on the result of the KS-test. In these cases, the ND function changes most intensively, while at the edges its change is not so pronounced. Therefore, when the outburst of mortality occurs in the middle segment of the change in the function, then the distribution is more likely to be non-normal than if the outburst occurs at the beginning or end of its segment. In addition, we found that when the sample size was increased by a factor of 2 (the number of deaths per day was multiplied by 2), in a situation where the number of repeated values in a number of LS increased by a factor of 2, the value of the D_n_ test statistic did not change, whereas the D-critical and p-value decreased in a regular way, which was expressed by the appearance of a non-normality estimate. Thus, the more individual flies participate in the experiment (which is certainly better in the general case), the more repeated values arise (since the maximum LS of the sample individuals is limited) and the greater the risk of defining the data distribution as non-normal increases. Thus, there is a bias and dependence of the KS-test result on the sample size: the smaller the sample size, the higher the probability of determining a normal distribution. This feature of the KS test was most clearly manifested when the sample was divided into intervals (the number of observation points was reduced), when we obtained a normal distribution for all cases studied.

Thus, having considered the mechanics of the KS test for single samples, we concluded that it cannot be applied to assess the strength of genetic background and to track the effect of mutations on ontogeny because of the strong influence of repeated values and outliers, i.e., variation in the distribution of data (violation of the linearity of variance). Consequently, we further considered only the SW test approach and interval-divided samples to analyse the LS data and predict biological patterns.

### 2.4. Qualitative and quantitative parameters of the Shapiro-Wilk test can be used to assess biological patterns in describing survival rates

As a model to investigate whether it is possible to detect biological patterns in interval samples using the SW test, we analysed the survival data of white mutants and the corresponding control lines with the same genetic background. Assessing the effect of the white gene on LS as an integral characteristic of ontogenesis is an important task, because, despite the fact that the history of the study of the white gene goes back more than 110 years (Morgan, 1910; Green, 2010)., the physiological significance of this gene in Drosophila, including its role in the control of LS, remains unclear (Jl et al., 2021; Sasaki et al., 2021; Xiao and Qiu, 2021; Mendoza-Grimau et al., 2024). We investigated the effects on LS of two different mutant alleles of the gene, w^67c23^ and w^1118^ (Table 1 and 2, respectively, columns 7,8 in each table). Only the range of intervals containing the data was used to assess the resulting distributions for normality. Marginal intervals (e.g., 0-4 and 65-69), if they contained a zero probability of mortality, were not used to calculate the W test statistic.

In the experiment with the allele it turned out that the distribution in 3 cases out of 4 was reversed (Table 1, column 8, cases of distribution reversal are highlighted in Bold Italics). Moreover, in the first 2 cases out of 3 the distribution changed from non-normal in the control to normal in the mutant, and in the last case — vice versa. Thus, the character of distribution of control data does not necessarily have to fulfil the criterion of normality and can be non-normal. The change in the character of the distribution is obviously related to the change in LS. Thus, in all cases where the nature of the distribution changed, the change in µ was more pronounced than in the situation where the distribution remained qualitatively the same (Table 1, column 4). The change in the nature of the distribution was consistently accompanied by changes in W values — when normality changed to non-normality, W values decreased and vice versa (Table 1, column 7). However, in the most recent case (w Oregon RC females and Oregon RC males), when the distribution in the mutants did not change, remaining normal, there was virtually no change in W. This correlates well with the fact that the survival curves of the mutant and control in this case are closest in shape and position to each other (Fig. 3D), whereas in all other cases they are significantly different (Fig. 3 A, B, C).

**Figure 3:**
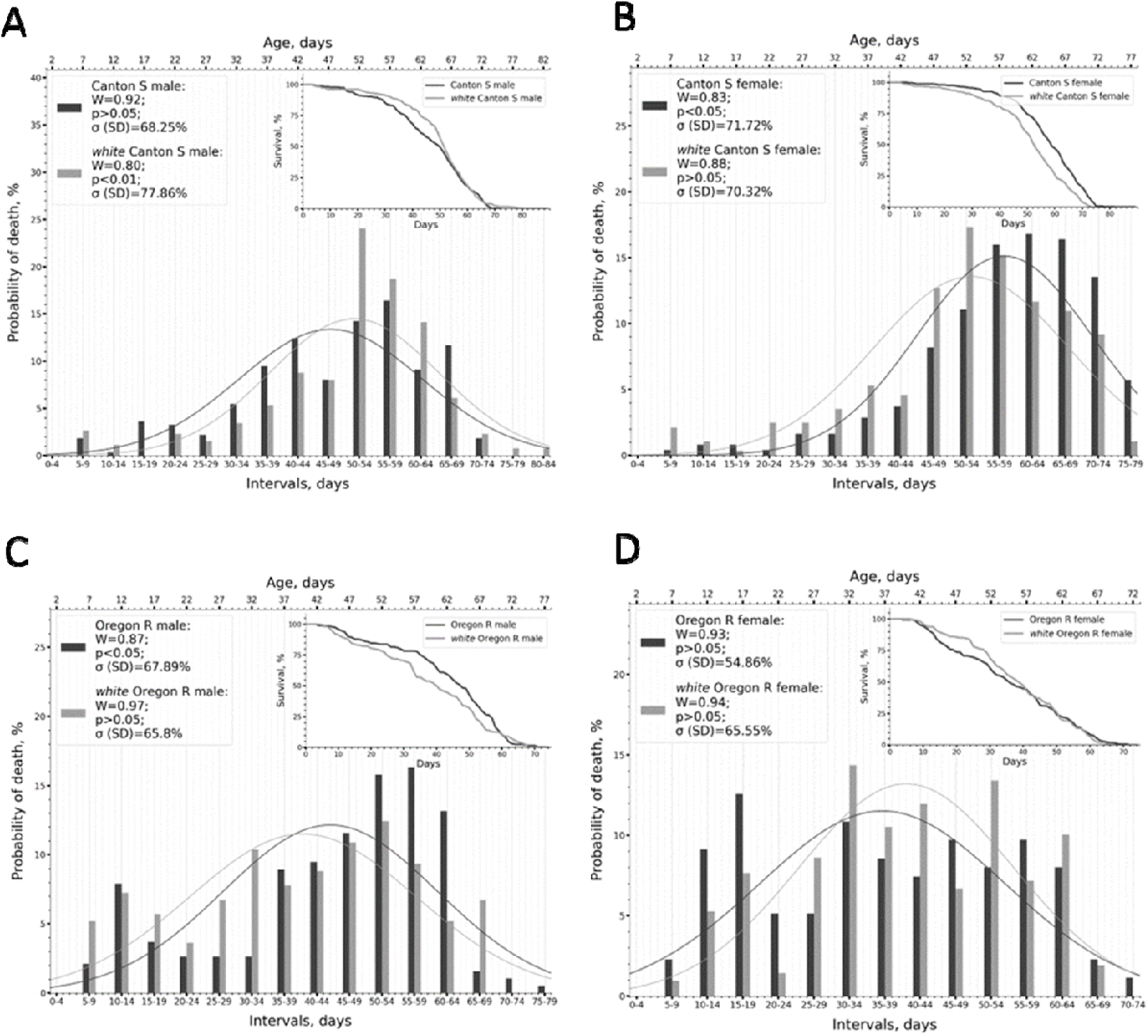
Experimental data with the allelew67c23. Experimental data distributions (bars) with phenotype frequencies obtained by dividing samples into 5-day intervals are shown. The names of the compared genotypes are indicated in the legend. Other notations are as in Figure 1E.

In the experiment with the allele^w1118^, it turned out that the distribution changed in the mutants, compared to the control, only for the Canton S genotype (Table 2, column 8, cases of distribution change are highlighted in Bold Italics). The distribution changed from normal in the control to non-normal in the mutant. The change in the pattern of distribution also as in the experiment with the allele^w67c23^ was probably related to the degree of change in LS. In the case of the Canton S genotype, the change in µ was generally more pronounced than in the case of the Oregon RC genotype (Table 2, column 4). The loss of normality of the distribution was accompanied by a consistent decrease in W values in the mutants (Table 2, column 7). The survival curves for mutant and control in the case of Canton S were strikingly different in shape and position relative to each other for both males and females (Figure 4 A, B). At the same time, for the Oregon RC genotype, there was no qualitative change in the distribution pattern upon introduction of the white mutation. However, changes in W values and p-value clearly indicated quantitative changes in probabilities (phenotype frequencies) within the sample. Thus, in males w^1118^ Oregon RS, the W value shifted towards normality from 0.77 in the control → to 0.84 in the mutant, and the p-value almost reached the normality boundary of 0.005 → 0.03 (normality threshold >0.05). Such changes, in our opinion, very subtly reflected the changes that occurred in the sampling structure after the introduction of the mutation, and describe them much better than survival curves: visually, survival curves in mutant males w Oregon RS and control Oregon RS males were almost indistinguishable in shape and position relative to each other (Fig. 4C). On the other hand, in the case of w^1118^ Oregon RS females, in contrast to control Oregon RS females, quantitative changes shifted toward abnormality. The W value from 0.91 in the control → decreased to 0.87 in the mutant, and the p-value became closer to the border of abnormality, 0.324 → 0.11 (abnormality threshold <0.05). At the same time, the survival curves of mutant females were shifted to the left, relative to the control, showing a decrease in LS, but the shape was not different from that of control females (Figure 4D).

**Figure 4:**
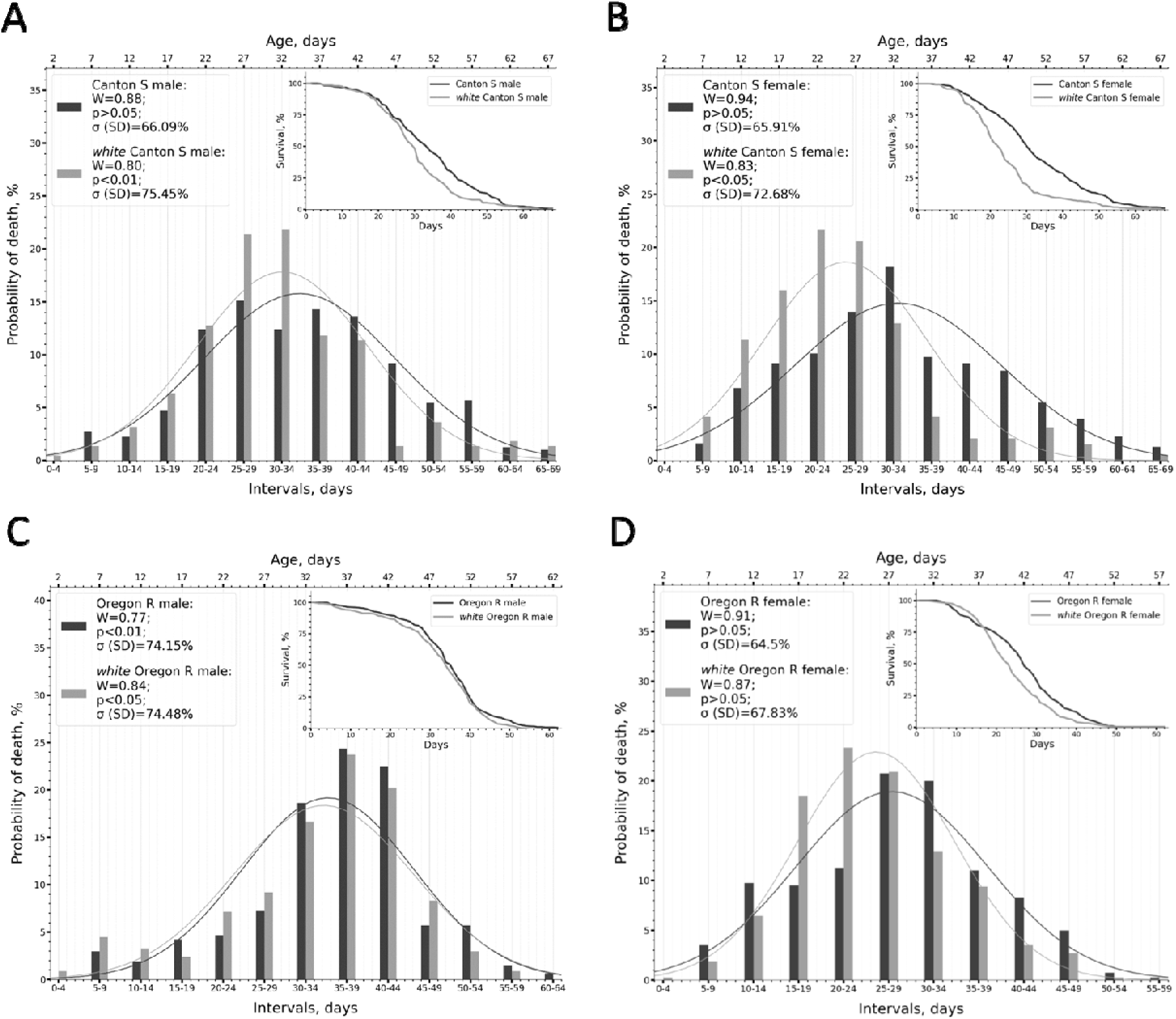
Experimental data with the w allele1118. Experimental data distributions (bars) with phenotype frequencies obtained by dividing samples into 5-day intervals are shown. The names of the compared genotypes are indicated in the legend. Other notations are as in Figure 1E.

Thus, the application of qualitative and quantitative normality criteria makes it possible to detect subtle differences in the sample structure even when the shape and/or position of the survival curves of the cohorts being compared repeat each other and are therefore hidden from the researcher’s attention.

### 2.5. Use of quantitative normality criteria to describe parameters of genetic health of a lineage

We determined that it is possible to use the normality criterion and its mathematical components (qualitative and quantitative) to generally describe i) the structure of the sample based on interval-disaggregated CV data, ii) assess what happens to the sample when interventions (introduction of the white mutation in our case) are made. At the same time, it remained unclear whether it was possible to directly relate LS to the parameters accompanying the normality assessment. To this end, we searched for correlations between the mean LS (µ), the standard deviation of µ (σ), and the SW test statistic (W), which reflects the normality of the distribution (the value of W varies from 0 to 1 and the closer it is to 1, the more likely it is that the data follow a normal distribution). The values of µ, as well as σ, were calculated from the LS data of all individuals in the sample, without dividing the data into intervals (the value of σ calculated from the intervals will be much wider than the real one). The results are presented in Table 3.

**Table 3:** Correlations between normality and parameters characterising the distribution of LS data.

| Correlated indicators | Correlation, $r$ | |
| --- | --- | --- |
|  | Experiment #1 w<sup>67c23</sup> | Experiment #2 w<sup>1118</sup> |
| $\mu$ (days) and $W$ | -0.78 | -0.09 |
| $\mu$ (days) and $\sigma$ (days) | -0.82 | 0.21 |
| $\sigma$ (days) and $W$ | 0.77 | 0.64 |
| CV* and $W$ | 0.80 | 0.54 |

\* CV was obtained by dividing the value of $\sigma$ (days) by $\mu$ .

It was found that there was a fairly strong correlation between µ and W in the case of the experiment with the allele, but no correlation in the case of the experiment with the allele (Table 3). The values of µ and σ were also not correlated with each other, but in both experiments, there was a clear positive correlation between the values of σ and W, as well as between the coefficient of variation (CV) (which is essentially σ normalised by µ) and W (Table 3). The presence of this correlation obviously follows from the µ criterion W itself, where the numerator of W, based on formula #15, reflects the “consistency” of the experimental data with the assumed normal distribution, and the denominator represents the variant σ — the variance. Thus, W is a measure of normality, normalised, in effect, by the size of the variability.

The observed direct correlation between W and σ deserves a closer look. To this end, we expressed σ for each sample not in days, as in Tables 1 and 2, but in %, indicating how many per cent of flies fell within the ±1σ interval and how many per cent of flies did not fall within this interval (how many remained) (Table 4). In a standard normal distribution, the number of individuals falling within the ±1σ interval ≈ 68.2%.

**Table 4:** Percentage of individuals falling and not falling within the ±1σ interval.

| Genotypes | Sex | Experiment #1 w<sup>67c23</sup> |  | Experiment #2 w<sup>1118</sup> |  |
| --- | --- | --- | --- | --- | --- |
| | | % of individuals<br>in the range $\pm 1\sigma$ | % of individuals<br>in the range $\pm 1\sigma$ | % of individuals<br>in the range $\pm 1\sigma$ | % of individuals<br>in the range $\pm 1\sigma$ |
| Canton S | M | 68.25 | 31.75 | 66.09 | 33.91 |
|  | F | 71.72 | 28.28 | 65.91 | 34.09 |
| *white* Canton S | M | 77.86 | 22.14 | 75.45 | 24.55 |
|  | F | 70.32 | 29.68 | 72.68 | 27.32 |
| Oregon RC | M | 67.89 | 32.11 | 74.15 | 25.85 |
|  | F | 54.86 | 45.14 | 64.50 | 35.50 |
| *white* Oregon RC | M | 65.80 | 34.20 | 74.48 | 25.52 |
|  | F | 65.55 | 34.45 | 67.83 | 32.17 |

Then we expressed the dependence of W on σ and on p-value on the graphs. It turned out that the normality criterion W is i) inversely related to the number of individuals falling within the ±1σ interval — the larger the value of W (i.e. the more normal the distribution), the smaller the number of dead individuals falling within the ±1σ interval (Fig. 5 A,B) and ii) directly related to the number of flies not falling within the ±1σ interval — the larger the value of W, the larger the number of individuals falling outside the ±1σ interval, i.e. the centre of the distribution (Fig. 5 C,D). This means that the more individuals fall outside the ±1σ interval, the wider the spread of LS values in days, which is what we found when looking for correlations.

**Figure 5.**
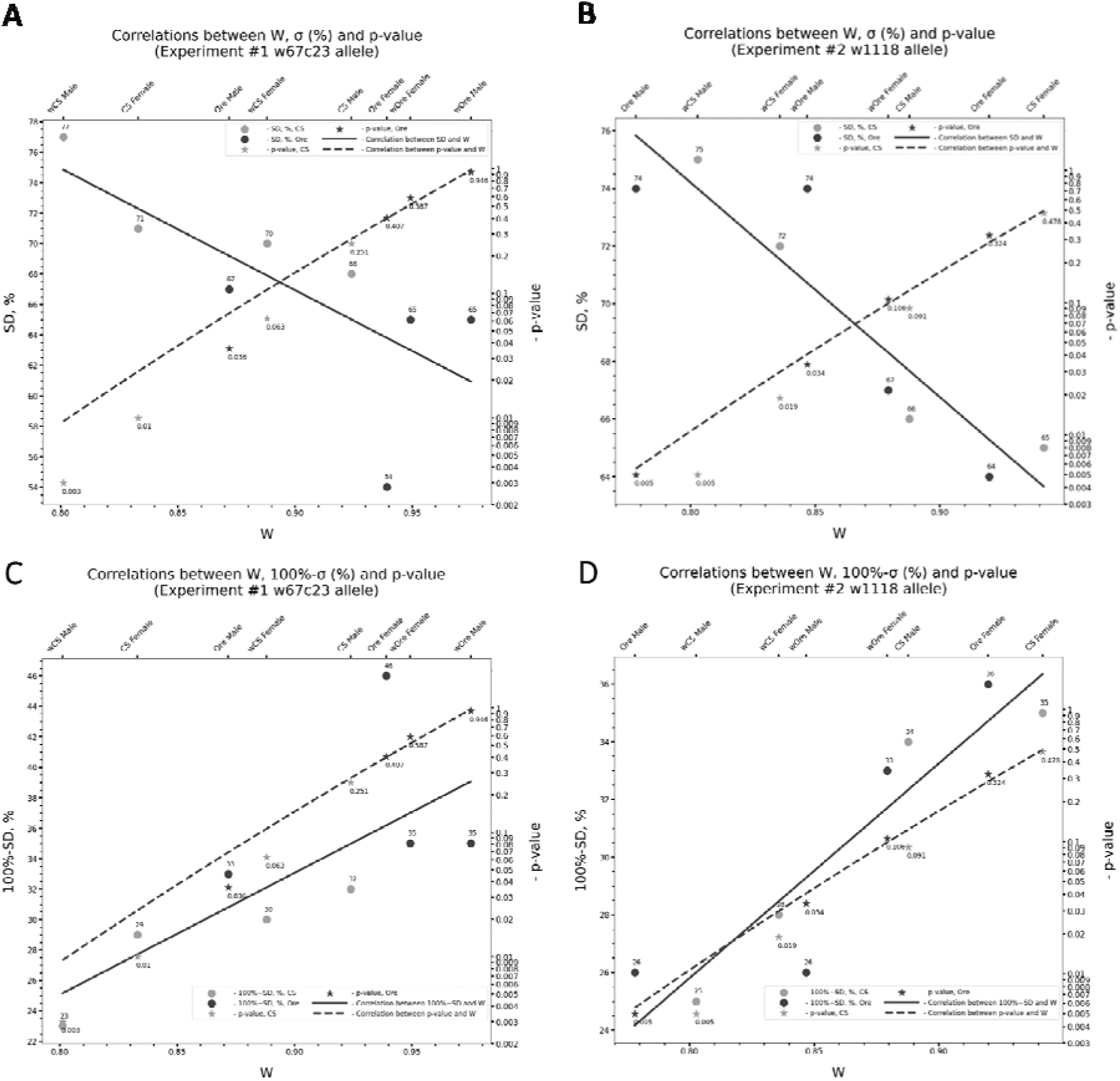
Plots of the dependence of W values on: (i) the number of individuals (in %) falling within the ±1σ interval (Y1 axis) and p-value (Y2 axis) for experiment #1 (allele w^67c23^) (A) and for experiment #2 (allele w^1118^) (B), (iii) numbers of individuals (in %) that did not fall within the ±1σ interval (Y1 axis) and from the p-value (Y2 axis) for experiment #1 (allele w^67c23^) (C) and for experiment #2 (allele w^1118^) (D). Captions on the X2 axis are genotype names: CS — Canton S, Ore — Oregon RC, w — *white*⍰. Numbers next to circles denote % of individuals from Table 4 for a given genotype, numbers next to asterisks denote p-values from Tables 1,2. The colour code of genotypes is indicated in the legend.

How can the dependence of W on σ be related to the LS values of the samples? A careful consideration of the dependencies indicated in the graphs in Figure 5 suggests that it is possible to classify genotypes according to the strength of genetic background on the basis of two indicators: normality criterion and µ (Table 5). Then: i) lines with increased LS would be characterised by a non-normal distribution due to the presence of increased late survival, combined with the absence or presence of increased early mortality. A double absence of both late survival and increased early mortality is also possible. Lines with these characteristics could be assigned an overall grade/rank of 3. Their distribution would be skewed to the right and their survival curves would be convex (e.g. Figure 3B), ii) lines with intermediate LS would be the most abundant, have a normal distribution and be characterised by the presence of increased early mortality combined with the absence of increased late survival. There may also be variants where early mortality is not elevated and elevated late survival is present or absent. Lines with normal distributions can be assigned an overall grade/rank of 2. Their distribution will be in the centre and the survival curves will be flat (e.g. Figure 3D), iii) lines with reduced LS will have a non-normal distribution and will be characterised by the presence of increased early mortality and the absence of increased late survival. It is also possible that there is no increased early mortality but no increased late survival. Such lines can be assigned an overall class/rank of 1. Their distribution would be shifted to the left and the survival curves would be concave inwards (e.g., Figure 4B).

**Table 5.** Classification of genotypes based on µ values and normality parameters.

| 1 | 2 | 3 | 6 | 7 | 8 | 9 | 10 |
| --- | --- | --- | --- | --- | --- | --- | --- |
| Genotype | Sex | $\mu$, days | Distribution characteristics* | | Increased early mortality** | Increased late survival** | Lifespan class (rank) |
|  |  |  | W | Distribution* |  |  |  |
| Experiment #1 $w^{67c23}$ | | | | | | | |
| Canton S | M | 47.9 | 0.92 | norm. | - | - | 2 |
|  | F | 58.3 | 0.83 | not normal. | - | + | 3 |
| $w^{67c23}$ Canton S | M | 50.7 | 0.80 | not normal. | - | + | 3 |
|  | F | 52.2 | 0.88 | norm. | - | + | 2 |
| Oregon RC | M | 44.6 | 0.87 | not normal. | + | + | 3 |
|  | F | 37.2 | 0.93 | norm. | + | - | 2 |
| $w^{67c23}$ Oregon RC | M | 39.4 | 0.97 | norm. | + | - | 2 |
|  | F | 39.4 | 0.94 | norm. | + | - | 2 |
| Experiment #2 $w^{1118}$ | | | | | | | |
| Canton S | M | 34.8 | 0.88 | norm. | - | + | 2 |
|  | F | 33.1 | 0.94 | norm. | + | + | 2 |
| $w^{1118}$ Canton S | M | 31.6 | 0.80 | not normal. | - | - | 1 |
|  | F | 25.2 | 0.83 | not normal. | + | - | 1 |
| Oregon RC | M | 35.2 | 0.77 | not normal. | - | - | 3 |
|  | F | 27.9 | 0.91 | norm. | + | - | 2 |
| $w^{1118}$ Oregon RC | M | 33.7 | 0.84 | not normal. | - | - | 3 |
|  | F | 25.1 | 0.87 | norm. | + | - | 2 |

\* — according to SW test calculated on data divided into intervals, \*\* — subjective assessment based on a review of graphs.

For each class, it is possible to calculate its mean LS. We summed µ values in males and females of control and mutant lines in each experiment (sum of 4 genotypes). Class 3, using the data in Table 5, was characterised by µ=44.5 days, class 2 by µ=38.99 days, and class 1 by µ=28.4 days.

Summing up the ranks of individual lines belonging to the same genotype allowed us to infer its strength. This complements the estimation based on µ values. According to the analysis in experiment #1 with the w allele (experiment #1), the Canton S genotype is strong (sum of ranks 10), whereas the Oregon RC genotype is weak (sum of ranks 9). An experiment with the w allele (experiment #2) performed with the same lines 12 years later showed that the strong genetic background of Canton S had converted to a weak genotype (rank sum of 6), whereas the Oregon RC genotype was unchanged (rank sum of 10). Apparently, in the Canton S line, when collection flies are kept in small populations (usually 2 tubes from which flies are mobilised into new tubes once a month to maintain the collection), allelic variants with weakly deleterious effects emerged, accumulated and stabilised due to genetic drift over a period of 12 years (Lynch et al., 1995; Latter, 1998; Gilligan et al., 2005). The reduced strength of the Canton S genetic background is clearly visible from the ratio of µ values between genotypes (Canton S vs. Oregon RC) calculated for experiment #1 and #2. Thus, the ratio between Canton S (µ=52.27 days) and Oregon RC (µ=40.15) genotypes for experiment #1 is 1.3, while for experiment #2 it is 1.02 (Canton S µ=31.17 days, Oregon RC µ=30.7). Thus, although experiments #1 was performed at a slightly lower temperature, which generally lengthens Drosophila LS (Flatt, 2020; Shaposhnikov et al., 2022). The degradation of the Canton S genotype is very apparent in the LS ratio between genotypes in each experiment.

Returning to the relationship between variability (σ) and LS (µ), we can conclude that a narrowing of σ, accompanied by the emergence of a non-normal distribution, will mean the appearance of long-lived and short-lived phenotypes in the sample, respectively, relative to the original normal distribution. We have expressed similar ideas in our previous work, suggesting that the change in LS occurs by changing the frequencies of occurrence of short- and long-lived phenotypes within a sample/population/cohort (Bylino et al., 2024) We hypothesised that in order to shift the values of the distribution of the LS trait to the right (towards increasing LS), variability on the left must be reduced and increased on the right. An alternative hypothetical approach, as we observed, might be to reduce the variability of the trait on both sides of the distribution (i.e., reduce the norm of the trait response) and then shift the centre of the trait distribution to the right, using several strategies aimed at stabilising the occurrence of different types of variability (Bylino et al., 2024) In this paper, having analysed the distributions of real samples, we conclude that changes in the frequencies of short- and long-lived phenotypes in one sample (experimental) relative to another (control), e.g. when ontogenesis is destabilised by mutation (genetic intervention) are observed across the entire frequency range (see, for example, the selected examples in Figures 3B,C and 4D), i.e. across the entire distribution. Thus, a shift of the centre of the distribution to the right (an increase in LS) is likely to be accompanied by a two-way change in frequencies both to the left and to the right of the centre of the distribution. Hence, unilateral changes in variability are apparently impossible. However, verification of this assumption requires additional study of distribution skewness and kurtosis for samples with non-normal distribution and will be considered in future work.

As for the decrease in variability to the left and right of µ and the shift of the centre of distribution to the right (increase in LS) without changing the level of variability, we see that the introduction of a mutation that increases LS (allele w^67c23^) results in a narrowing of the σ range and the appearance of new phenotypes in the entire frequency range. At the same time, the frequencies of phenotypes of the original distribution continue to be present in the sample. Hence, it seems that moving the narrowed distribution to the right is impossible without increasing the variability of the LS trait itself. Thus, the only way to increase the LS of the sample is to expand the variability of the LS trait.

Thus, the main results of the paper can be summarised in the following conclusions:

1. Partitioning the survival curve data into intervals (solution to problem 1) allows: i) minimising non-linearities in the variance of daily mortality, i.e. getting rid of dips/outliers in mortality values while maintaining the general trend of the distribution, ii) visualising and efficiently analysing the data distributions for normality by the Shapiro-Wilk test using convenient tables of critical values, iii) the Kolmogorov-Smirnov test for single samples cannot be used to assess normality as for data partitioned into indices.
2. The application of qualitative and quantitative criteria of normality and distribution can be used as reinforcing and complementary ways of analysing survival to describe biological patterns, the strength of genetic background and the impact of interventions (solution to problem 2). It allows: i) to see differences between samples indistinguishable by other means (estimation of mean LS, visual inspection of survival curves), ii) to rank lines according to the strength of genetic background. However, there is no correlation between LS and quantitative normality criteria, but a correlation between normality and standard deviation of LS. More research will be needed to clarify how quantitative normality criteria can be linked and used in assessing the strength of genetic background.

## Conclusion

In this paper, we tried to address two problems: the problem of representing the structure of the survival curve data in a representative way (problem 1) and the use of normality criteria to assess the strength of the genetic background of the lines and the LS interventions performed (problem 2).

The proposed approach of partitioning the survival (survival) curve data into intervals solves the problem of adequately estimating the sampling structure, which is not at all clear when considering curves (problem 1). This new layer of information about sampling structure is extremely useful for understanding what is happening in an experiment and allows visualisation in a simple way of comparing an experimental distribution with a control distribution or two experimental distributions. Additional formalisation is achieved by overlaying the interval data representation of bell-shaped curves with a normal distribution calculated from the full sample data.

Regarding the application of normality criteria to assess the strength of genetic background, we expected to find a clear relationship between the mean LS and the quantitative normality score. We expected that W would show a dependence on µ and, varying from 0 to 1, would serve as a measure of the strength of the genetic background of a line. As it turned out, this was not the case, and instead we found a correlation between W and σ. Thus, W obviously cannot act as an independent criterion for assessing the strength of genetic background and cannot be used as a mathematical approximation/analogue to µ or, for example, the maximum LS of a line. Thus, the use of normality criteria does not completely solve the problem of mathematical evaluation of the strength of genetic background and the influence of the conducted interventions on LS, but, for the time being, appears to be only an additional tool for analysis. However, even now, as we have shown, when applied with care to the data, the researcher has a new analysis tool that helps to detect changes in biological processes with mathematical precision, in addition to the classic tests for differences in median, mean and maximum LS.

More research will be needed to understand how normality criteria can be properly applied in the best way in survival analyses. New approaches are needed to assess the strength of genetic background and the impact of genetic interventions.

## Author Contributions

Conceptualization, O.V.B.; Formal analysis, K.A.V., A.A.S., O.V.B.; Investigation, K.A.V., O.V.B.; Methodology, K.A.V., O.V.B.; Resources, Y.V.S., A.A.S.; Project administration, O.V.B.; Supervision, O.V.B.; Funding acquisition, O.V.B.; Writing and editing, K.A.V., O.V.B. All authors have read and agreed to the published version of the manuscript.

## Institutional Review Board Statement

Not applicable.

## Informed Consent Statement

Not applicable.

## Data Availability Statement

Data are contained within the article.

## Acknowledgments

The authors thank the Science for Life Extension Foundation, represented by its president Mikhail Meltzer and philanthropist Alexey Khozyaykin and his company Glavny Masterplace, for supporting the research. The authors are also grateful to Alexander Trofimov for his invaluable advice on mathematics.

